# An effector XopL enhances ethylene biosynthesis to promote *Xanthomonas fragariae* infection in strawberry

**DOI:** 10.1101/2024.04.11.589074

**Authors:** Xiaolin Cai, Wenyao Zhang, Jinnan Luo, Wei Li, Rui-hong Chen, Xiang-nan Xu, Ying-qiang Wen, Jia-yue Feng

## Abstract

*Xanthomonas fragariae* (*Xaf*) is the cause for strawberry crown dry cavity rot and strawberry leaf angular spots. Despite having a long evolutionary history with strawberries, the plant-pathogen connection is poorly understood. Pathogenicity for the majority of plant pathogens is mostly dependent on the type-III secretion system, which introduces virulence type III effectors (T3Es) into eukaryotic hosts cells. For most of these T3Es, the subcellular targets are yet unclear. Here, We used the yeast-two-hybrid (Y2H) technique to construct an interaction network of strawberry-*Xaf* T3Es. Multiple T3Es were discovered to converge onto the strawberry 1-aminocyclopropane-1-carboxylic acid oxidases (ACOs), which are the last rate-limited step in the production of ethylene. We then concentrated on the connection between XopL and FveACO9. Strawberry plants that overexpressed XopL accumulated higher levels of ethylene and exhibited more severe *Xaf* infection. XopL boosted ethylene production by stabilizing the accumulation of FveACO9 protein and enhancing ACO enzyme activity. Additionally, strawberries treated with ACC or overexpressing FveACO9 were particularly vulnerable to *Xaf* infection. On the other hand, pre-treatment with α-aminoinoisobutyric acid (AIB), an ACO inhibitor, effectively reduced *Xaf* infection. Our research indicates that *Xaf* utilizes a distinct approach to regulate the ethylene production of host plants in order to promote infection.

## Introduction

*Xanthomonas fragariae (Xaf)*, a bacterial pathogen highly specific to wild and cultivated strawberry, often causes angular leaf spots (ALS), and causes a huge production and economic loss in strawberry. Although strawberry ALS was reported first in the United States in Minnesota in 1960 (Kennedy, 1960), and since then it has spread worldwide, only a few is known regarding the epidemiology (Bestfleisch et al., 2015; Wang et al., 2023) and limited control options exist (Haack et al., 2019; Wang & Turechek, 2016). What’s noteworthy is that almost all strawberry germplasm was found to be either somewhat resistant or susceptible to this infection (Maas et al., 2000, 2002; Roach et al., 2016; Wei et al., 2024). Earlier research have given insight into the interaction of strawberry-*Xaf* using RNA-Seq (Gétaz et al., 2020; Puławska et al., 2020). Most importantly, the genetic information for around 74 Xaf strains was gradually made public (Gétaz et al., 2017; Henry & Leveau, 2016; Vandroemme et al., 2013), including two new strains YL19 and YLX21, which resulted in more severe symptoms in strawberry plants, namely crown dry cavity rot, excluding ALS (Li et al., 2021; Liang et al., 2023). So far, the fact that so little is known about the intricate process of interaction between strawberries and *Xaf* may be connected to the challenges in overcoming the technical barriers associated with *Xaf* genetic modification (Vandroemme et al., 2013), as well as the lack of resistant germplasm. Hence, developing an alternative novel strategy to discover the virulent factors of *Xaf* and reveal the susceptibility mechanism of strawberry to *Xaf* is essential for the prevention of this pathogen.

Pathogens often deploy the highly conserved type-III secretion system (T3SS) to transfer virulence proteins, known as type III effectors (T3Es), into eukaryotic host cells to invade their hosts and influence plant responses to favor infection (Büttner & Bonas, 2003; Dangl & Jones, 2019; Duxbury et al., 2016). There are 25 T3Es identified in the draft genome of *Xaf* strain LMG25863 (Vandroemme et al., 2013), 47 T3Es in *Xaf* strain Fap21 identified by machine-learning (Wagner et al., 2023), 33 T3Es in *Xaf* strain YL19 in our previous study based on homolog blast (Wei et al., 2022), yet most of these T3Es remain functionally uncharacterized. Finding the proteins that these putative pathogenic effectors target on their host will be essential to assigning roles to them. Honestly, numerous investigations have shown that T3Es are likely to concentrate onto a small number of intricately linked cellular hubs in order to impede pathogen fitness and effectively inhibit host defense. For example, in *Pseudomonas syringae-Arabidopsis thaliana*, two studies have identified plant protein TCP14 as the extensive target of T3Es by systematically effector-host protein interactions at the effectome-scale using a systematic Y2H (González-Fuente et al., 2020; Weßling et al., 2014). In addition, many independent experiments also identified common targets by multiple unrelated T3Es, such as *Arabidopsis* BAK1 are associated with AvrPto, AvrPtoB, and HopF2 of *P.syringae* (Shan et al., 2008; Zhou et al., 2014), four additional *P.syringae* T3Es (AvrRpt2, AvrRpm1, AvrB, and HopF2) directly targets and modifies RIN4, a plant immune regulator, to suppress PTI signaling (Axtell & Staskawicz, 2003; Mackey et al., 2002, 2003; Wilton et al., 2010). In fungi, a variety of *Phytophthora* Avr3a-like effectors target cinnamoyl alcohol dehydrogenases in the bid to adversely influence plant immunity. (Li et al., 2019). Not surprisingly, different host proteins also could be targeted by a single effector (González-Fuente et al., 2020), as in the case that the soybean Type 2 GmACSs are targeted and destabilized by the *Phytophthora sojae* RXLR effector Avh238 in order to inhibit ethylene production and encourage infection(Yang et al., 2019). Hence, T3Es could be used as probes to identify important components hub of resistance signaling pathways in plant.

Ethylene (ET) is a gaseous hormone that plays a variety of physiological, morphological, and stress-related roles in plants (Khan et al., 2017). In the ethylene biosynthetic process, S-adenosylmethione is transformed into 1-aminocyclopropane-1-carboxylic acid (ACC) by ACC synthase (ACS), which is subsequently converted into ethylene by ACC oxidase (ACO), which is encoded by a multigene family in higher plants (Houben & Van de Poel, 2019). Although ethylene has a well-established involvement in the plant defense response, its interactions with other and sometimes opposing plant immune responses make it a complex player (Li et al., 2023; Zhao et al., 2017; Zhao et al., 2022). Pathogen effectors could target ET biosynthesis and signaling pathways to compromise host immunity. For instance, *Xanthomonas euvesicatoria* (*Xe*) deploys XopD targets tomato transcription factor SlERF4 to directly repress ET biosynthesis and ET-stimulated defense (Kim et al., 2013). *P. syringae* HopAF1 targets the Yang cycle proteins MTN1 and MTN2 of *Arabidopsis* to interrupt ethylene accumulation to suppress PAMP-induced defense response during bacterial infection(Washington et al., 2016). However, exogenous ethylene treatment leads to enhanced plants resistance against different pathogens (Núñez-Pastrana et al., 2011; Gaige et al., 2010). In addition, the complete pathogenicity of rice dwarf virus, *Xanthomonas campestris* pv *vesicatoria*, *P. syringae pv glycinea*, and *Verticillium dahlia* on their hosts requires the generation of ethylene by the host (van Loon et al., 2006; Weingart et al., 2001; Zhao et al., 2017). Therefore, the functions of ethylene in plant defense vary depending on the context, taking into account the various pathogen threats and ways in which they interact with other defensive signals.

Through high-throughput yeast two-hybrid coupled with next-generation sequencing (Y2H-Seq), we discovered in this work that strawberry FveACOs are the target hub of *Xaf* T3Es. The interaction between XopL and FveACO9 was then investigated, and it was shown that overexpressing XopL increases ethylene production and increases the susceptibility of strawberry plants to disease. To encourage the production of ethylene, XopL increases the enzymatic activity of ACO and stabilizes the FveACO9 protein. Strawberry ethylene production and sensitivity to *Xaf* are increased by overexpression of FveACO9. Bacterial populations and symptoms of disease could be considerably reduced by exogenous ethylene inhibitors. In summary, the results of this investigation provide light on a distinct mechanism elucidating the role of ethylene as a negative regulator in strawberry defense against a bacterial disease.

## 2 Results

### 2.1 Strawberry ACC oxidases are the target hub of multiple *X.fragariae* type III effectors

To determine the target hub of *Xaf* effectors in strawberry, twenty-two putative T3Es were selected for identifying host targets via Y2H-Seq (Table S1). Before the screening, the subcellular location of each effector was identified by transient expression in *N.benthamiana* (Fig. S1). Five T3Es (XopL, XopN, XopAE, XopB, and XopF1) are associated with membrane-localization, which were selected to use the split-ubiquitin-based membrane yeast two-hybrid (MYTH) system, other 17 T3Es using nuclear Y2H system (nY2H) (Fig. S1, 2). In addition, among the 22 bait constructs, the full length of four effectors (XopD, XopAE, XopB, XopF1) showed autoactivation in the Y2H system. However, the truncated type of XopD and XopF1 lossed the self-activation function could be used for interactor mining (Figure S2), while the truncated variants of XopAE and XopB have no positive clones in the Y2H were discarded. Thus, the remaining twenty effectors were used as baits in a prey library composed of *F.vesca* cv. ‘Hawaii 4’ complementary DNA (cDNA), which has been challenged with *Xaf* YL19 before RNA extraction. The final data consisted of 1089 interactions between 20 T3E baits and 741 strawberry preys (Table S2; Fig. S3a). 42% of the identified proteins were associated with more than one of the tested candidate effectors (Fig. S3b), on average, a candidate effector associated with 55 host proteins, ranging from 2 for XopX2 to 204 for XopL (Fig.S3c), suggesting a high rate of promiscuity. Among them, the FveACO9 was targeted by seven T3Es (XopL, XopH, XopC1, XopE5, AvrBs1, XopX1, XopR) in the Y2H screening (Table S2; Fig. 1a), which was determined as the target hub. In adddition, XopL was associated with other 2 of 31 putative strawberry ACOs (FveACO16, FveACO18) in the interactor library (Fig. 1a, S4). We further examined that XopL also interacted with other two ACOs (FveACO14, FveACO21) in the thirty-one successful cloned FveACOs in yeast (Fig. 1b). Collectively, these results revealed that strawberry ACOs are possibly the target hub of multiple *Xaf* T3Es and XopL may play a key role in function to FveACOs.

**Figure 1.**
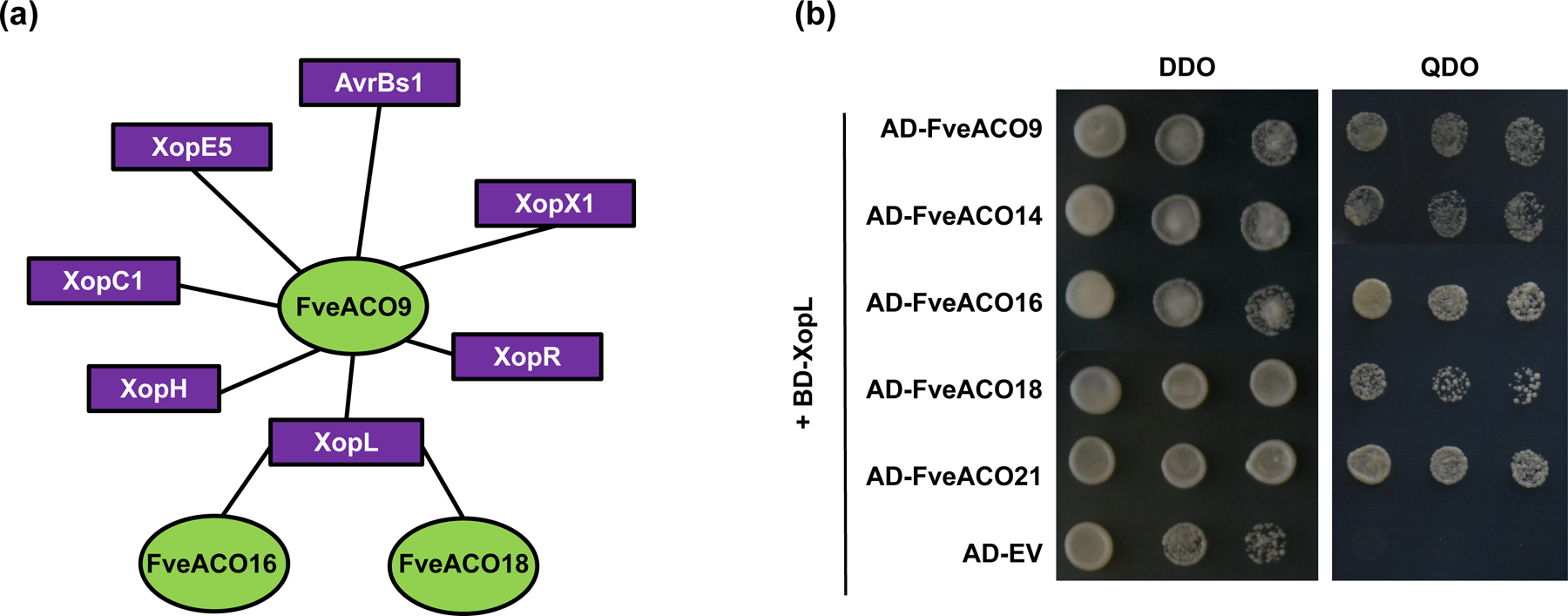
Strawberry ACC oxidases are the target hub of multiple putative *X.fragariae* type III effectors. (a) Multiple ACC oxidases of strawberry (*Fragaria vesca*) were positive preys of *Xaf* T3Es in the high-throughput Y2H screening. (b) XopL interacts with FveACO9, FveACO14, FveACO16, FveACO18, and FveACO21 in yeast, pGBKT7 (BD)-XopL and pGADT7 (AD)-empty co-transformation in Y2H-Gold yeast as negative control. Plasmids of the bait and prey pairs were co-transformed into yeast cells and selected on double dropout (DDO, SD/-Trp/-Leu) or quadruple dropout (QDO, SD/-Trp/-Leu/-Ade/-His) for 3-5 days.

### 2.2 XopL is secreted effector and interacts with FveACO9 *in planta*

To study the role of effectors in the function of strawberry ACOs and their impact on plant immunity. XopL and FveACO9 from the above yeast screening were chosen for further investigation. The split GFP method was used to investigate the translocation of XopL into host cells (Park et al., 2017). A nuclear localization signal (NLS) was attached to the N-terminal of GFP_1-10_ (named NLS-GFP_1-10_) and transiently expressed by agrobacterium for 24 hours in strawberry leaves. Subsequently, the leaves were inoculated with *Xaf* YL19 carrying XopL-GFP_11_. Figure 2(a) demonstrates that there is no GFP fluorescence in wild type leaves infected with *Xaf* YL19(XopL-GFP_11_) or in NLS-GFP_1-10_-expressing strawberry leaves. However, green fluorescence was seen in the nucleus when Xaf YL19(XopL-GFP_11_) infected NLS-GFP_1-10_ transiently expressed strawberry leaves. The data indicate that XopL possibly be translocated into host cells.

**Figure 2.**
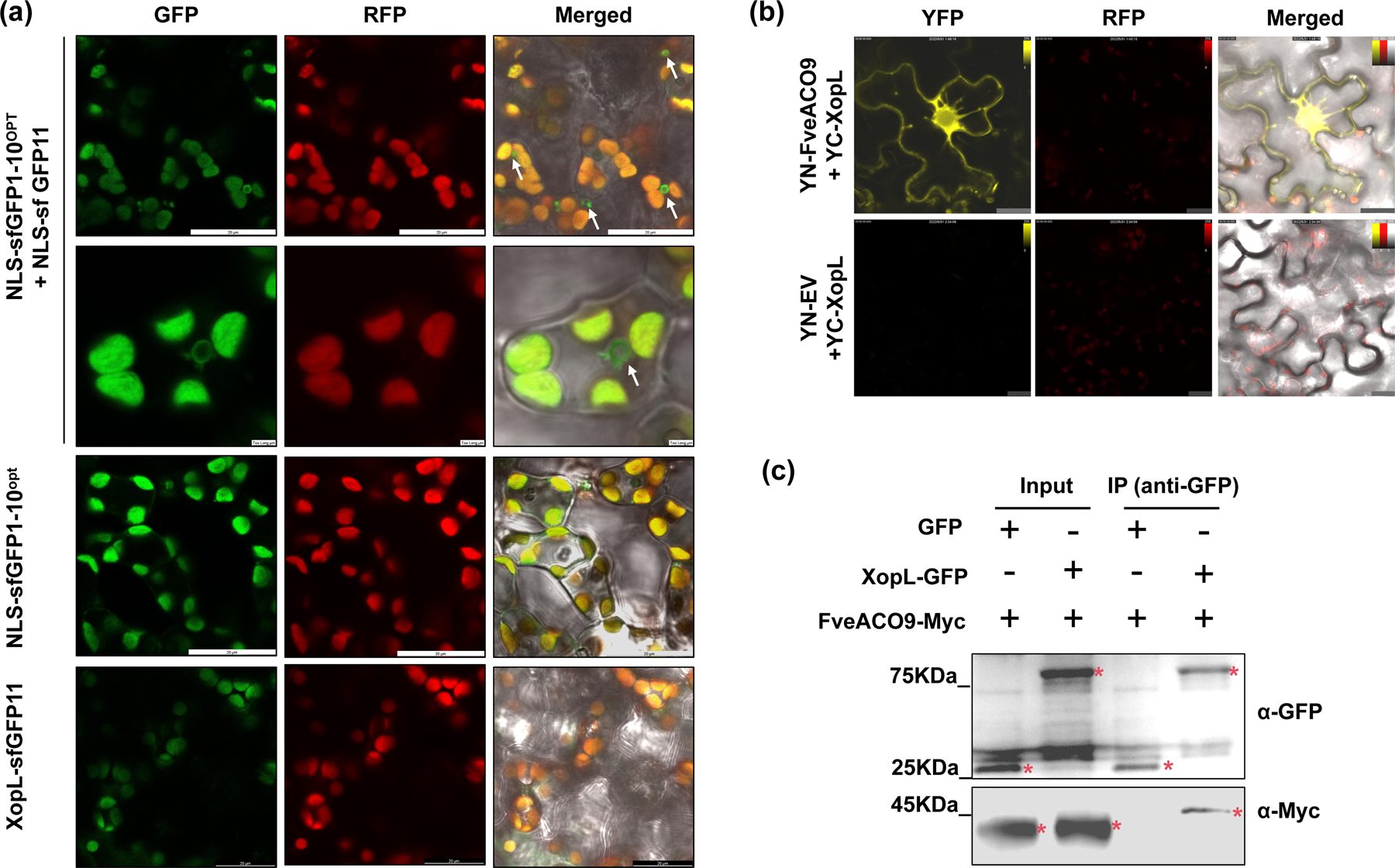
XopL translocated into host cells and interacts with FveACO9 in planta. (a) XopL translocation assay using split GFP system. The transient expressed NLS-Myc-sfGFP1-10^OPT^ strawberry leaves were inoculated by Xaf YL19(XopL-sfGFP11-HA). Images were obtained at 3 days post-inoculation. Bars scale 20 μm. The white arrow indicates XopL-sfGFP11-HA and NLS-Myc-GFP1-10^OPT^ colocalization signaling. (b-c) XopL interacts with strawberry FveACO9 in bimolecular fluorescence complementation (BiFC) (b), and Co-immunoprecipitation (Co-IP) (c) assays. The proteins used for Co-IP were derived from N.benthamiana co-expressed XopL-GFP with FveACO9-Myc. Total proteins were then subjected to incubation with GFP-Trap_A beads and anti-Myc (α-Myc) immune blotting of the output was used to identify the co-immunoprecipitation of XopL-GFP with FveACOs-Myc. The red asterisk indicates the expected protein and its molecular masses are XopL-GFP (81 KDa), GFP (26.8KDa), and FveACO9-Myc (41KDa). +, protein expressed; -, vector control.

**Figure 3.**
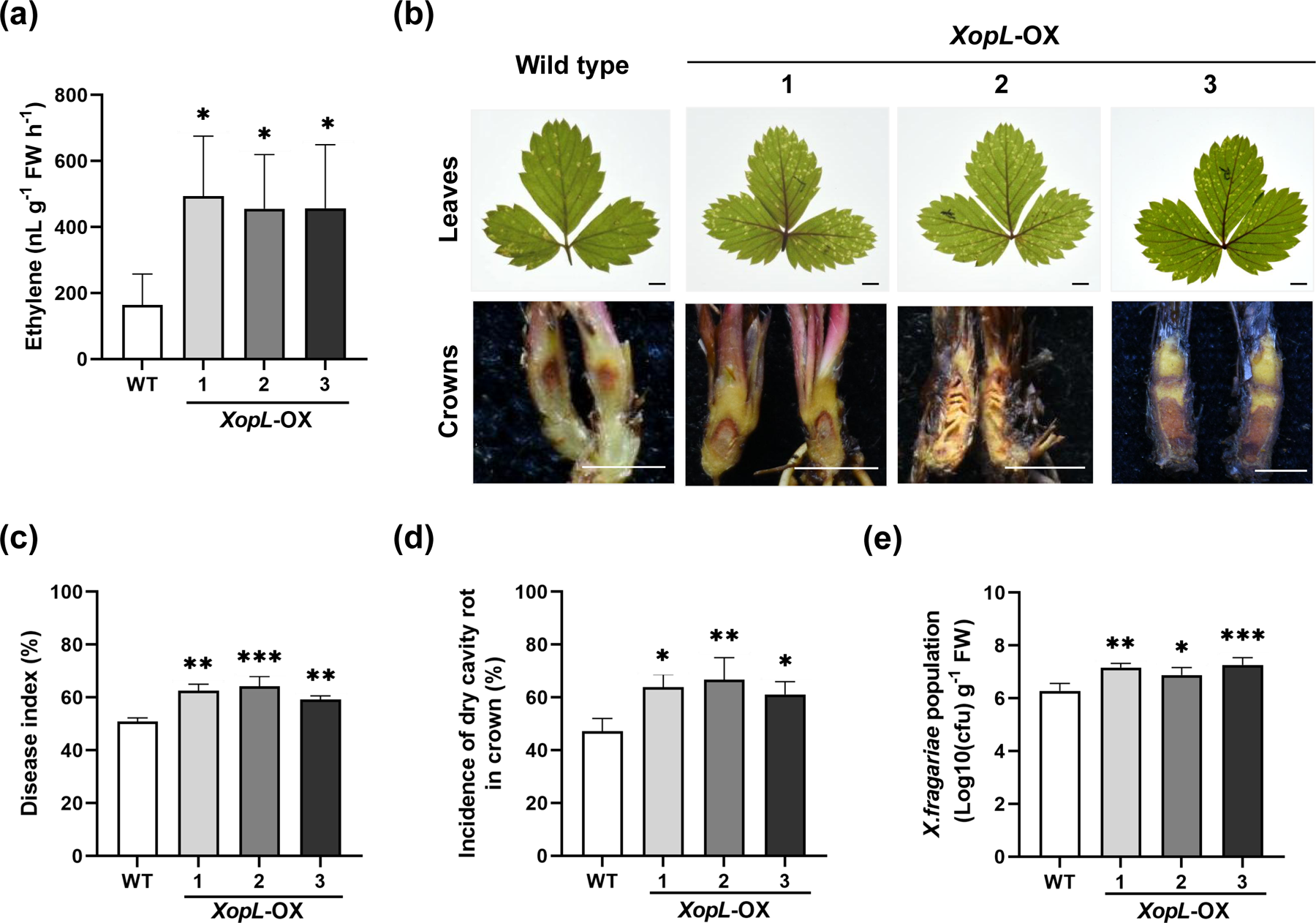
Overexpression of XopL enhances ethylene accumulation and *X.fragariae* susceptibility in strawberry. (a) Ethylene accumulation of the wild type and *XopL*-OX strawberry lines. Data are mean ± SD (n=4). (b) Disease symptoms in leaves and crowns of wild type and *XopL*-OX strawberry lines were photographed at 7, and 45 days post inoculation (dpi), respectively; (c-e) Leaf disease index (c), and the incidence of crown dry cavity rot in inoculated strawberry plants (d), and bacterial population in wild type and *XopL*-OX strawberry lines (e). Data are mean ± SD (n=3). Asterisks indicate significant differences between WT and *XopL*-OX strawberry lines by Two-way ANOVA (Dunnett’s multiple comparisons test). *, P<0.05; **,P<0.001, ***, P<0.001.

Furthermore, bimolecular fluorescence complementation (BiFC) assay was carried out to verify whether XopL interacts with FveACO9 in planta. N-terminal nYFP-tagged XopL and N-terminal cYFP-tagged FveACO9 co-expressed in *N.benthamiana*, the empty cYFP coexpressed with nYFP-XopL as negative control. The results showed that no YFP fluorescence signal in the negative control, while nYFP-XopL co-expressed with cYFP-FveACO9 in *N.benthamiana* exhibited bright YFP fluorescence signal, suggesting that XopL interacts with FveACO9 *in planta* (Fig. 2d).

We further confirmed that XopL interacts with FveACO9 in *N.benthamiana* extracts using co-immunopreciptation assay. *N.benthamiana* leaves were hand-infiltrated with the suspension of A.tumefaciens coexpressing XopL-GFP and FveACO9-Myc (Fig. 2e). A solubilized total protein extract was isolated from leaves 48 h post infiltration, and then FveACO9-Myc was pulled-down using GFP agarose beads. Protein gel blot analysis shows that FveACO9-Myc copurified with XopL-GFP but not GFP. These results demonstrate that XopL physically associates with FveACO9 in plant extracts.

### 2.3 Overexpression of XopL enhances ethylene accumulation and disease susceptibility in strawberry

Considering ACO is the final rate-limiting step of ethylene biosynthesis and targeted by XopL. We were interested in investigating the effect of XopL on the ethylene biosynthesis in strawberry. Thus, three independent GFP-fused XopL transgenic lines (named *XopL*-OX) were generated and XopL-GFP protein was confirmed by confocal microscopy and Western blot (Figs. S5a, b). No visible phenotypic differences were observed between the wild-type and *XopL*-OX lines in normal growing conditions (Fig. S5b). Nevertheless, *XopL*-OX strawberry lines easily suffered early senescence in natural drought conditions (Figs. S6a, b). Measurement of ethylene has been shown a significantly elevated level in *XopL*-OX lines than the wild type (Fig. 2a), but similarly not at the expense of ACC, the ethylene precursor, which showed a mild change in *XopL*-OX lines in comparison to the wild type (Fig. S5c). These results indicated that XopL induces leaf senescence may by enhancing ethylene levels in strawberry.

To investigate whether XopL plays a pathogenic role in *Xaf* YL19 infection, inoculation tests using *Xaf* YL19 showed that more severe infection symptoms with a larger area of angular spot in leaves at 7 dpi (Fig. 2b-c) and higher percentage of crown dry cavity rot in the inoculated plants at 45 dpi than the WT plants (Figs. 2b, d). Consistently, a significantly higher bacterial replication in *XopL*-OX leaves than in the wild type (Fig. 2e). Collectively, these results showed that overexpression of *XopL* compromised strawberry defense to *Xaf* YL19, and that was possibly associated with the increased ethylene level

### 2.4 XopL promotes FveACO9-catalyzed ethylene biosynthesis and activates ethylene signaling pathways

To determine FveACO9 whether catalyze ethylene production and the effect of XopL on the activity of FveACO9 *in planta*. FveACO9 alone or with XopL were co-expressed in *N.benthamiana* via agroinfiltration. The leaves were collected at 24 hpi, followed by measurement of ET by gas chromatography. *N. benthamiana* leaves expressing FveACO9 showed significantly higher ethylene production than those GFP-expressing control leaves (Fig. 4a). However, XopL alone did not markedly alter the ET production, but significantly promoted the ET production catalyzed by FveACO9 (Fig. 4a). These results indicated that FveACO9 is possibly a functional ACO enzyme that could convert ACC into ethylene, and XopL promoted FveACO9-catalyzed ethylene biosynthesis.

**Figure 4.**
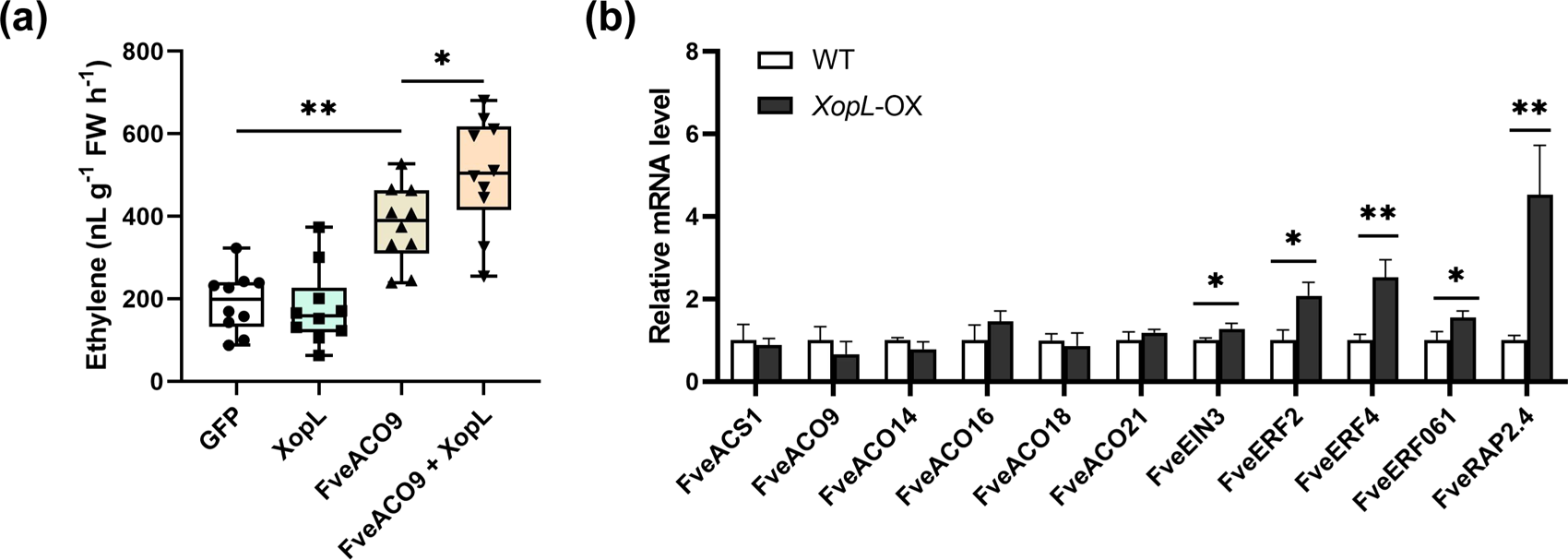
XopL promotes FveACO9-catalyzed ethylene production *in planta*. (a) Measurement of ethylene production at 48 hours after transient overexpression of GFP-fused FveACO9 alone or co-expression with XopL in *N.benthamiana*, the GFP and XopL-GFP as negative controls. Data are mean ± SD (n=10). (b) The transcript level of ethylene biosynthesis and signaling transduction-related genes in the wild type and *XopL*-OX strawberry. Data are mean ± SD (n=3). Asterisks indicate significant differences between WT and *XopL*-OX strawberry lines by Student’s *t*-test . *, *P*<0.05; **,*P*<0.001.

Furthermore, we examined whether XopL regulates ET biosynthesis at the transcriptional level, the mRNA abundance of seven ET biosynthesis genes (*FveACS1, FveACO9, FveACO14, FveACO16, FveACO18, FveACO21*) and five ET signal transduction genes (*FveEIN3*, *FveERF2*, *FveERF4*, *FveERF061*, and *FveRAP2.4*) were determined. Compared to the wild type, there were no significant differences in ET biosynthesis genes in *XopL*-OX lines. However, *FveEIN3*, *FveERF2*, *FveERF4*, *FveERF061*, and *FveRAP2.4* were significantly up-regulated in *XopL*-OX lines (Fig. 4b). These results demonstrated that XopL greatly triggers the ET signal pathways but has little impact on the accumulation of mRNAs involved in ET biosynthesis.

### 2.5 XopL enhances protein accumulation of FveACO9 and ACO enzyme activity

Because of a significantly elevated ethylene level but less cost of ACC and a WT transcription level of *FveACO9* in the presence of XopL, we speculated the stability of FveACO9 may be enhanced by XopL. Hence, the luciferase-tagged FveACO9 was co-expressed with XopL-GFP in *N.benthamiana* for analysis of protein stabilization, and three negative control pairs were designed as shown in Figure 4(a). The LUC/XopL-GFP and LUC/GFP co-expression, respectively, led to similar but lower luminescence intensity than FveACO9-LUC co-expression with or without XopL in *N.benthamiana*, while FveACO9-LUC coexpressed with XopL-GFP had more strong luminescence intensity than it coexpressed with GFP (Figs. 5a, b). The results indicated that the protein level of FveACO9 could be stabilized in the presence of XopL. Also, immunoblotting analysis supported the result (Fig. 5c).

**Figure 5.**
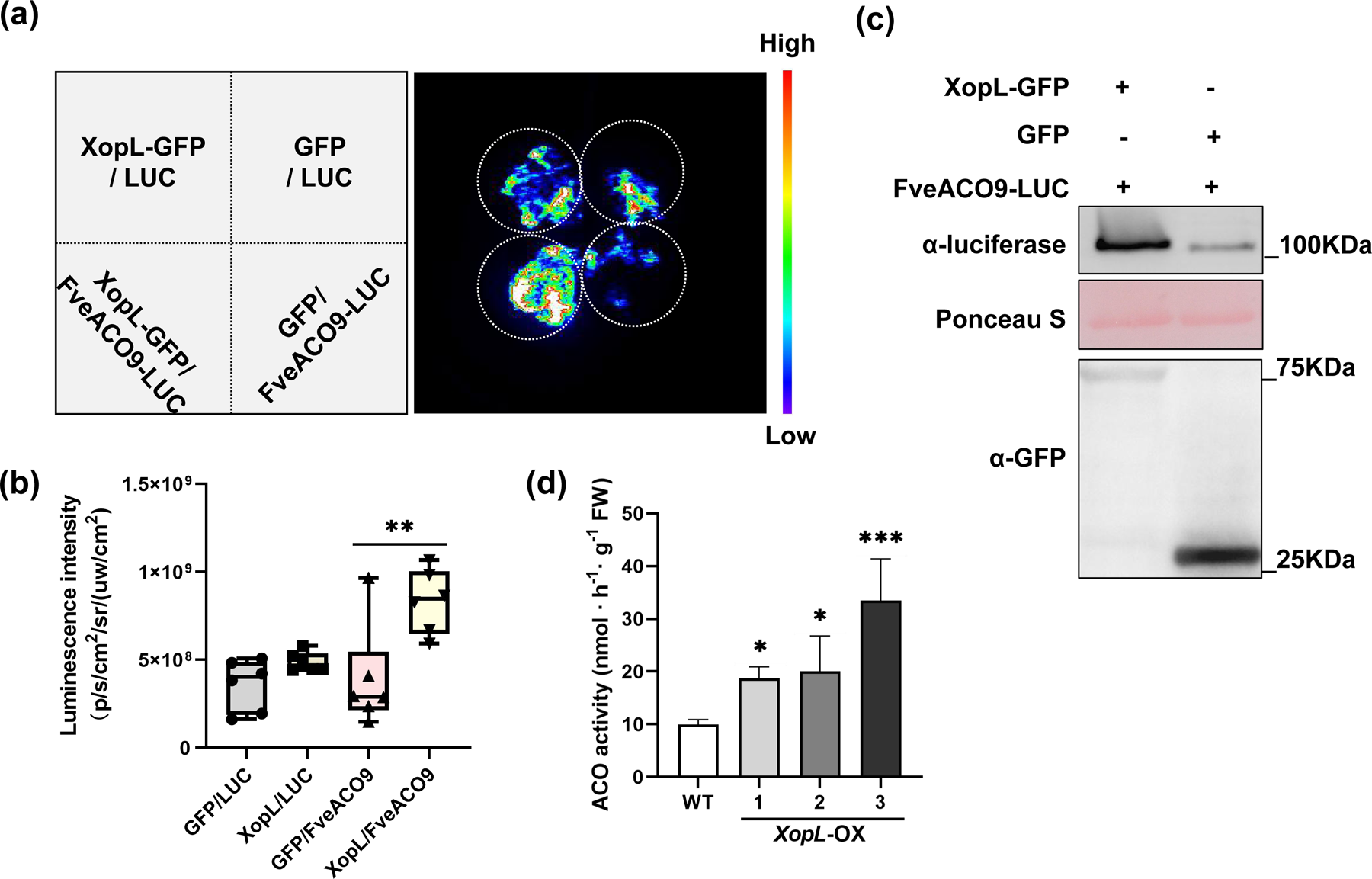
XopL promotes protein accumulation of FveACO9 and ACOs enzyme activity . (a-b) The representative images of bio-fluorescence (a), and measurement of bio-fluorescence intensity in *N.benthamiana*, data are mean ± SD (n=6), in which XopL-GFP co-expressed with FvACO9-LUC, FveACO9-LUC/GFP, LUC/XopL-GFP, and LUC/GFP as negative controls (b). Asterisks indicate significant differences between FvACO9-LUC/XopL-GFP and FveACO9-LUC/GFPW ( Student’s *t* test). **, *P*<0.01.(c) Protein abundance of FveACO9-LUC was determined using anti-luciferase. The Ponceau S staining shows equal protein loading. Protein molecular masses are XopL-GFP (81 KDa), GFP (26.8KDa) and FveACO9-LUC (100KDa),. + protein expressed; -, vector control. (d) ACO enzymatic activity in the wild type and *XopL*-OX strawberry. Data are mean±SD (n=5). Asterisks indicate significant differences between WT and *XopL*-OX strawberry lines by Two-way ANOVA (Dunnett’s multiple comparisons test). *, *P*<0.05; ***, *P*<0.001.

Furthermore, the total enzymatic activity of FveACOs in *XopL*-OX lines was detected. As expected, the three *XopL*-OX strawberry lines presented significantly higher ACO enzyme activity than the wild type (Fig. 6d). Collectively, XopL promotes ethylene production and may not only stabilize FveACO9 protein levels but also enhance ACO enzyme activity.

**Figure 6.**
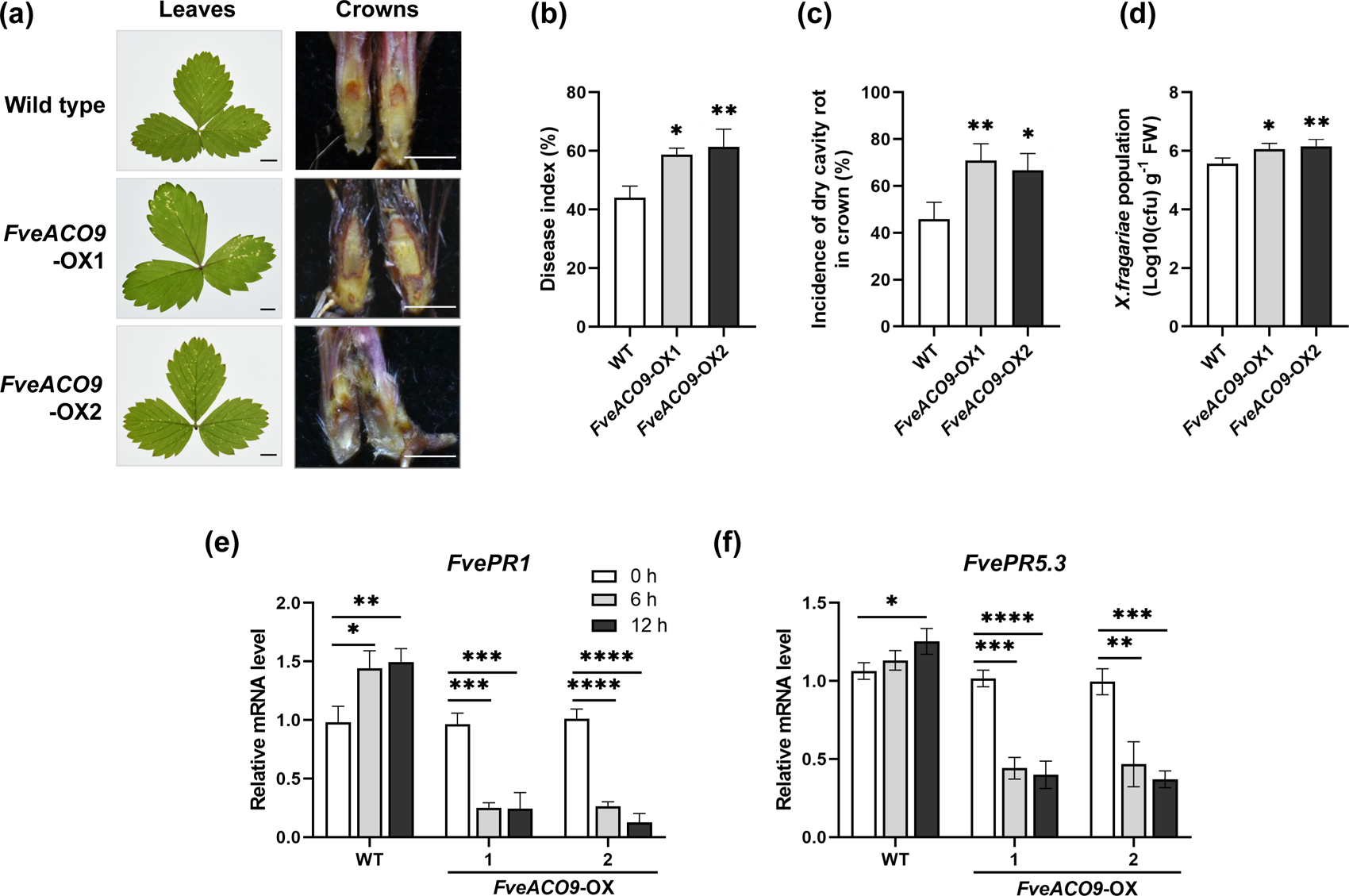
Overexpression of FveACO9 caused strawberries more susceptible to *X. fragariae* YL19. (a) Representative symptoms of the wild type and *FveACO9*-OX strawberry lines with *Xaf* YL19 inoculation, the symptom images of leaves and crowns were taken at 7 dpi = 50 mm. (b) Leaf disease index (%) and (c) incidence of crown dry cavity rot (%) was recorded at 7 dpi and 45 dpi, respectively. Data are mean ± SD (n=3). (d) The bacterial population of the wild type and *FveACO9*-OX strawberry lines at 7 dpi; (e-f) Relative mRNA level of *FvePR1* and *FvePR5.3* in wild type and *FveACO9*-OX strawberry lines at 0, 6, and 12 h after inoculation with *Xaf YL19* were analyzed using quantitative RT-PCR. Data are mean ± SD (n=3). The asterisks on the top of each column indicate a significant difference between WT and *FveACO9*-OX strawberry lines by Two-way ANOVA (Dunnett’s multiple comparisons test). *, *P*<0.05; **, *P*<0.01, ***, *P*<0.001, ****, *P*<0.0001.

### 2.6 FveACO9 mediated strawberry more susceptibility to *X.fragaria* YL19

To further ensure the function of FveACO9 in strawberry during *Xaf* YL19 infection, the luciferase-fused FveACO9 was introduced into the strawberry to generate *FveACO9*-overexpression (named *FveACO9*-OX) lines (Fig. S7a). We also produced transgenic strawberry plants carrying the *FveACO9* RNAi construct (knockdown) (Fig. S7c). Two *FveACO9*-OX lines were used to further analyzed, and the protein level of FveACO9-LUC in strawberry plants was confirmed by Western blot (Fig.S7b). The relative expression of *FveACO9* were about 21.9%-40.7% down-regulated in RNAi lines were confirmed by qRT-PCR (Fig. S7d). The *FveACO9*-OX and RNAi lines did not exhibit any morphological phenotypes difference with the wild type (Figs. S7a, c). The ethylene contents increased in *FveACO9*-OX lines but no significant difference in the *FveACO9-*RNAi lines, compared to the wild type plants (Fig. S7e). These results further proved that FveACO9 does promote ethylene production in strawberry.

To investigate whether *Xaf* infection, bacterium accumulation, and the host response is affected by ethylene contents in the *FveACO9*-OX lines, we inoculated fifteen seedlings from each line (WT, OX1, OX2) with *Xaf* YL19 and observed the resulting disease symptoms. At 7 dpi, those two *FveACO9*-OX lines showed more severe angular spot symptoms and higher disease index in leaves (Fig.6a,b), as well as a higher percentage of crown dry cavity rot at 45 dpi (average at 63.8 %) than the wild type control plants (average at 47.2%). qRT-PCR analysis revealed that bacterial population was higher in the *FveACO9*-OX leaves than in the wild-type (Fig.6c). We also tested the resistance level of *FveACO9*-RNAi lines, but the results showed that no significant difference whatever in symptoms or bacterial population between RNAi lines and wild type (Fig. S8a-d). These results suggested that overexpression of *FveACO9* enhances *Xaf* infection whereas knockdown of only FveACO9 is not enough reduces *Xaf* infection.

To analyze the possible mechanism of FveACO9-mediated susceptibility to *Xaf* YL19, we determined the expression of two pathogen-related genes *FvePR1* and *FvePR5.3* after treatment with *Xaf* YL19. The expression level of both *FvePR1* and *FvePR5.3* exhibited a significant decrease in *FveACO9*-OX lines than the wild type control plants at 6 and 12 h post inoculation (Fig. 6e, f). Collectively, these results revealed that FveACO9 is possibly a negative regulator of strawberry resistance to *Xaf* YL19, which is related to the down-regulation of defense gene expression.

### 2.7 Ethylene accumulation increases disease susceptibility in strawberry

To measure strawberry ethylene evolution during *Xaf* YL19 infection, we inoculated *Xaf* YL19 in strawberry plants, and the control plants were treated with H_2_O. The results showed that ethylene of the infected strawberry leaves significantly quick released over a 96-h course of the infection compared to the control plants (Fig. 7a), whereas the level of ACC only significantly decreased at 6 hours post-inoculation (Fig. 7b). The results suggested that ethylene was induced, but may not at the expense of ACC, by *Xaf* YL19 infection.

**Figure 7.**
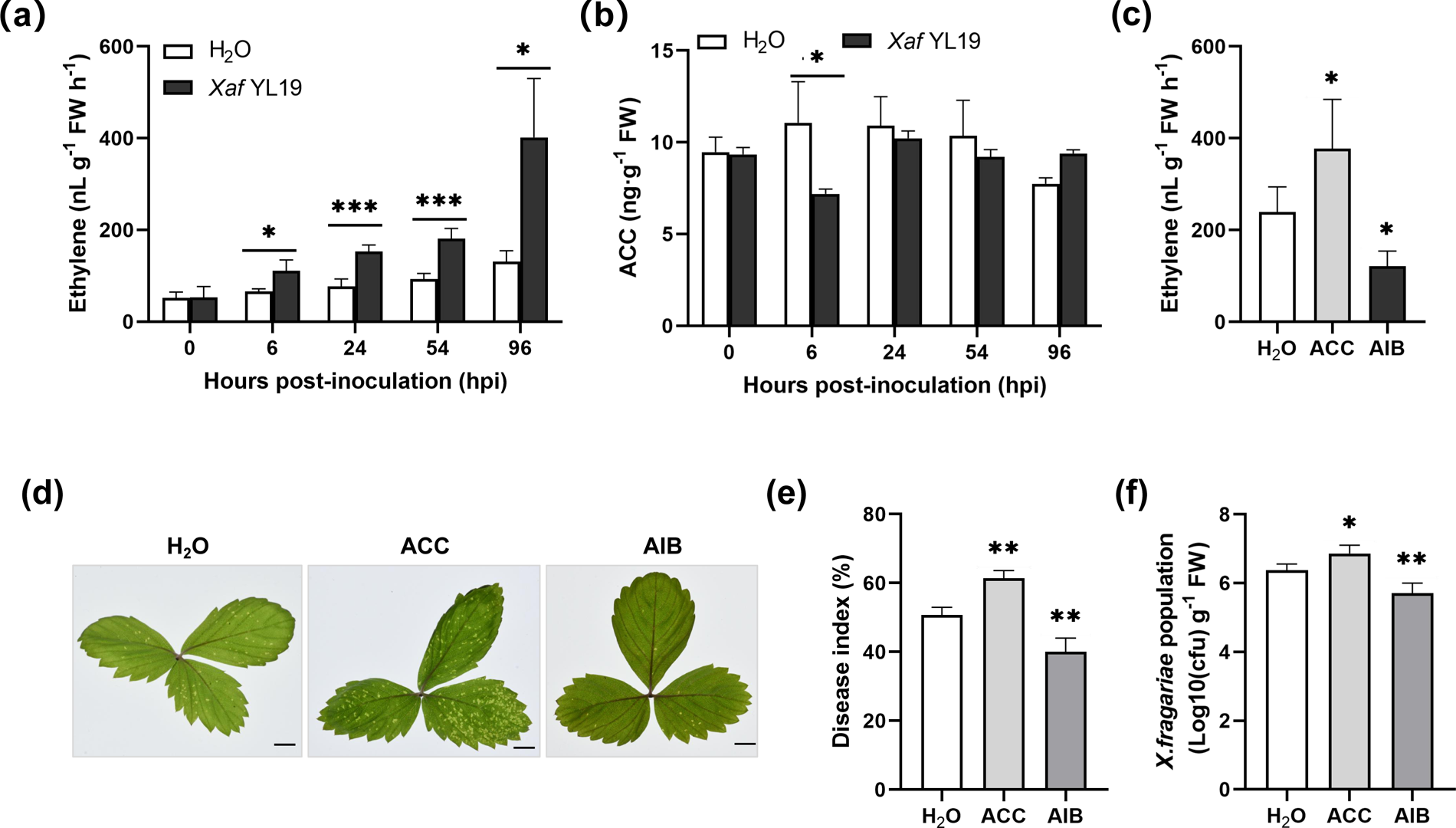
Ethylene is induced by *X.fragariae* infection and ethylene accumulation increases strawberry susceptibility. (a-b) The level of ethylene (a), and ACC (b) in strawberry plants at 0, 6, 24, 54, and 96 h post inoculation with *Xaf* YL19. Treatment with H_2_O as controls. Data are mean ± SD (n=3). Asterisks indicate significant differences between the treatments of H_2_O and *Xaf* YL19 (Student’s *t*-test *P* values, *, *P*<0.05; ***, *P*<0.001). (c) Ethylene level of strawberry plants after treatment with H_2_O, 30 μM 1-aminocyclopropane-1-carboxylic acid (ACC), or 50 μM α-aminoisobutyric acid (AIB). Data are mean ± SD (n=5). (d) Phenotypes of angular spots on strawberry leaves. Photographs representative of three independent experiments were taken at 7 d post-inoculation (dpi). Strawberry leaves were first sprayed with 30 μM ACC, 50 μM AIB, and H_2_O was used as the control. (f) *Xaf* YL19 population was quantified by quantitative polymerase chain reaction (qPCR) at 7 dpi. Data are mean ± SD (n=3). The asterisks on the top of each column indicate a significant difference between H_2_O and ACC or AIB treatment by Two-way ANOVA (Dunnett’s multiple comparisons test). *, *P*<0.05; **, *P*<0.01.

To further determine whether ethylene is involved in strawberry susceptibility to *Xaf* YL19 infection, we pretreated strawberry with 30 μM ACC, 50 μM AIB (α-aminoisobutyric acid, the ACOs inhibitor), or H_2_O as control for 12 h. Ethylene level was significantly increased or decreased by ACC or AIB treatment compared to the H_2_O-treated control, respectively (Fig. 7c), revealing that the ACC and AIB treatments worked as expected. Then, the pretreated plants inoculated with *Xaf* YL19 suspension. At 7 dpi, the ACC-treated plants exhibited greater susceptibility to *Xaf* infection, displaying more severe disease symptoms and bacterium accumulation than the *Xaf*-infected plants teated with H_2_O. However, the AIB-treated plants showed enhanced disease tolerance, as shown by less bacterial accumulation and milder disease symptoms, when compared with the *Xaf*-infected plants treated with H_2_O (Fig. 7d-f). Taken together, these data suggest that, ethylene is induced by *X.fragariae* infection and ethylene negatively regulates strawberry susceptibility to *Xaf*.

## 3 Discussion

In a previous study, we discovered that during *Xaf* YL19 infection, strawberry cultivars with moderate resistance showed lower ethylene levels than cultivars with high sensitivity (Wei et al., 2022). In this investigation, we verified that *Xaf*-induced ethylene accumulation occurs (Fig. 7a) and discovered by a Y2H screening that many T3Es are associated with the last essential enzyme of ethylene biosynthesis, ACOs, in strawberry plants. Additionally, using the interactor pair XopL and FveACO9 as an example, we presented multiple lines of evidence demonstrating that, as the straightforward model shown in Figure 8, XopL stabilizes FveACO9’s protein accumulation and initiates ACOs enzyme activity to increase ethylene production and ethylene signaling, thereby undermining the host resistance response and favoring pathogen colonization.

**Figure 8.**
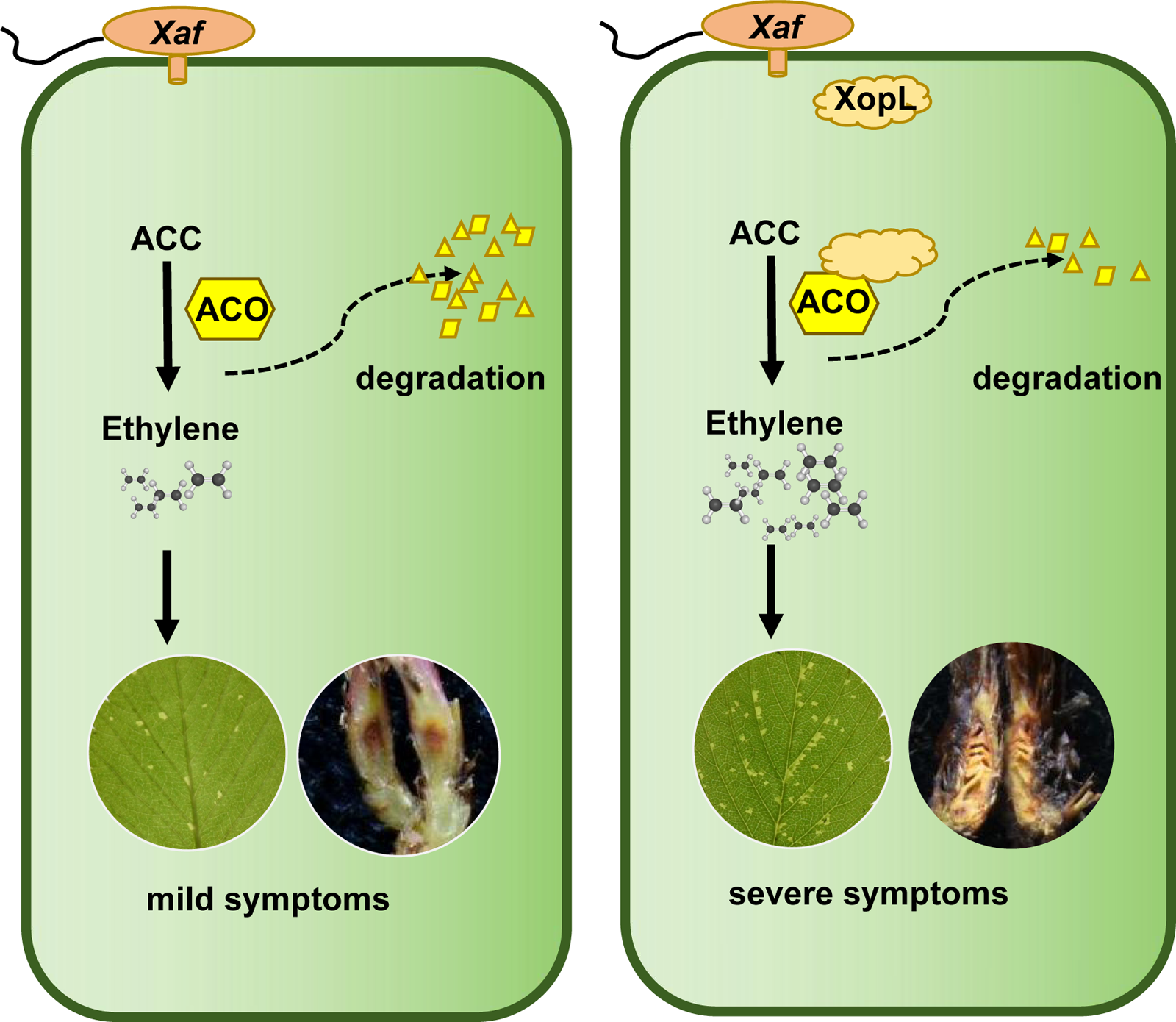
A schematic summary of XopL promotes *X.fragariae* YL19 virulence. 1-aminocyclopropane-1-carboxylate oxidases (ACOs) are the last rate-limiting step of ethylene biosynthesis, and ethylene plays a negative role in the response to *Xaf* infection. *Xaf* secretes the effector XopL that stabilizes ACO protein accumulation and triggers ACO enzyme activity to promote ethylene production and increase infection.

Phylogenetic analyses have focused on 31 putative ACOs in *F. vesca*, with 5 of these ACOs being targeted by XopL. However, few studies have used purified recombinant ACOs to characterize their functionalities, so it is unclear whether the putative FveACOs utilized in these analyses truly convert ACC into ethylene (Houben & Van de Poel, 2019). According to our results, *FveACO9* overexpression in *N. benthamiana* (Fig. 4a) does raise the amount of ethylene; the same result is supported by stable overexpression of *FveACO9* in strawberry plants (Fig. 3). Under many physiological conditions, a common approach to improve ethylene signaling seems to be the upregulation and/or stability of key enzymes in the ethylene production pathway, such as ACSs and ACOs. For instance, MPK6’s phosphorylation of ACS2 and ACS6 stabilized them in response to flg22 treatment, increasing ethylene synthesis and signaling(Liu & Zhang, 2004). Given these precedents, it may not seem surprising that FveACO9 gets stabilized to promote ethylene production by XopL to weaken defense response (Fig. 3c, 4b). It is intriguing that XopL contains nine LRR domains (Fig. S9), which are responsible for interacting with proteins(Kobe, 2001). In our Y2H screening, XopL exhibited the highest number of interactors and is associated with five FveACOs (Figs. S3c, 1a). These targets might potentially form hetero- or homo-dimerization with XopL’s assistance, hence enhancing the stability of FveACO9. Furthermore, the control of ACO enzyme stability and activity is connected to a number of posttranslation processes, including the S-nitrosylation of ACO4 in tomatoes by NO(Liu et al., 2023), and S-sulfhydration of ACO4 in Arabidopsis (Aroca et al., 2015). Through site-directed mutagenesis tests, two recent investigations have shown the significance of redox-specific cysteine changes to regulate ACO activity and structural stability(Fournier et al., 2019; Tachon et al., 2019). Therefore, it would be fascinating to see how XopL activates the ethylene-signaling pathway and increases protein accumulation and enzyme activity of FveACO9 to promote ethylene synthesis in further research.

Overexpression of both XopL and FveACO9 resulted in increased ethylene production along with impaired resistance to *Xaf* (Figs. 3, 6). However, FveACO9 silencing did not yield speculate results, showing that a WT level of resistance and ethylene was present in FveACO9-RNAi strawberry lines (Fig. 5). This could be easily explained by the fact that there are thirty-one potential functional ACO family members in strawberries, five of which may be enhancing ethylene evolution with the aid of XopL (Figs. S4, 1b). Therefore, silencing one member of the ACOs family may not be sufficient to significantly reduce ethylene production and improve remarkable resistance. However, because of the extremely varied nucleotide sequence in the *F. vesca* ACO family (Fig. S10), we were unable to overcome the technical obstacles to mute or knockout all FveACOs or just these five ACO interactors. As an alternative strategy, the application of exogenous ACO enzyme activity inhibitor AIB enhanced the level of resistance to *Xaf* (Fig. 7), giving a slight of ACOs or ethylene acts as a negative factor for strawberry resistance to *Xaf* (Fig. 7). One example of this was the interaction between tomatoes and *Salmonella*, where the tomato mutants with deficiencies in ethylene generation, perception, and signal transduction showed a significant reduction in Salmonella growth (Marvasi et al., 2014). Ethylene signalings are negatively related to plant defense responses (Fig. 6e, f), and to some extent, ethylene signaling possibly affects the expression of the unknown or known pathogen genes involved in the interactions with plants (Marvasi et al., 2014). Furthermore, XopL stimulated the ethylene signal pathway gene expression (Fig. 4c), which is in line with the finding that 1.51-fold up-regulation of ERF5 (FvH4_5g19800) was seen after *Xaf* infection(Gétaz et al., 2020), indicating an important role for ethylene signaling in *Xaf*-strawberry interaction. Thus, further studies will deepen our cognition about the function of ethylene and its pathway components in *Xaf*-strawberry interaction.

In conclusion, by inhibiting plant defensive mechanisms, the XopL targets and takes advantage of FveACO9, the negative regulator of plant innate immunity, to make pathogen invasion easier. It will also need further investigation to determine why ethylene helps the *Xaf* infection and to verify if ethylene has the same effects on strawberry infections caused by other bacteria. Furthermore, being the last stage of ethylene production, gene editing on ACOs largely lessens the impact on other pathways while potentially enhancing resistance.

## 3 Methods

### 3.1 Plasmid construction and plant transformation

For expression of GFP11-tagged XopL protein in *X. fragariae* YL19, The XopL gene from the genomic DNA of *Xaf* YL19 was amplified by polymerase chain reaction (PCR). The synthesized DNA fragments of GFP_11_ (Tsingke, China) were tagged to the C-terminal of XopL and subsequently ligated into pBBR1MCS5 using *EcoR*I and *BamH*I producing pBBR1MCS5(XopL-GFP_11_) was electro-transfers into the competence of *Xaf* YL19 and cultured in Nutrient Broth (NB) medium containing gentamycin at 25 ℃. N-terminal of GFP_1-10_ tagged nuclear location signal (NLS) was cloned into pCAMBIA2300 under the control of the 35S promoter, creating pCAMBIA2300(NLS-GFP_1-10_) and electro-transferred into Agrobacterium GV3101, and culture in LB medium containing kanamycin, gentamycin and rifampcin at 28 ℃. For plant transformation, the coding region of XopL was amplified from genomic DNA prepared from *X.fragariae* YL19. Full length cDNA for FveACO9 was amplified by PCR from cDNA pools prepared from leaves of strawberry (*Fragariae vesca* Hawaii 4). The full length of XopL and FveACO9 fused with GFP or luciferase, respectively, then cloned into pCAMBIA2300 vector to control expression with the cauliflower mosaic virus (CaMV) 35S promoter. For RNAi vector, a 289-bp (base pair) fragment of FveACO9 was amplified, and cloned into pRNAi vector, the same 289-bp fragment was subcloned into the vector in the inverse orientation. The corresponding recombination vectors were transformed into *A. tumefaciens* (GV3101) for plant transformation as previously described (Ma et al., 2023). Transgenic plants were identified by PCR and immunoblotting. Primers are described in Supplementary Table 3.

### 3.2 Effector translocation assay

A method based on previous research (Park et al., 2017) with minor revisions was used for effector translocation assay in strawberry plants. Briefly, the Agrobacterium strains GV3101 with NLS-GFP_1-10_ were streaked onto an LB agar plate with 200 μM acetosyringone (AS) and left to incubate at 28℃ for two days. After that, the bacteria were transferred to an infiltration buffer that contained 1/2 MS, 1% sucrose, 200 μM AS, and 0.01% Silwet L-77. The bacteria were washed two or three times, and the OD_600_ was diluted to approximately 0.5. The two-month-old strawberry seedlings were vaccum-agroinfiltrated. After infiltration, the plants were allowed to air dry for one hour before being placed in darkness for twenty-four hours. The plants were then inoculated with *Xaf* YL19(XopL-GFP_11_) for two days. Confocal microscopy was used to monitor fluorescence signaling at 488 nm excitation.

### 3.3 Pathology assays

*Xanthomonas fragariae* strain YL19-GFP was preserved in -80℃ from previous study used for pathology assay(Wang et al., 2023). YL19-GFP was streaked onto NB medium containing kanamycin (50 μg ml^-1^) for 5 days at 25 ℃. The whole strawberry plants were spraying-inoculated with YL19-GFP suspensions (1×10^8^ cfu ml^-1^) . The images of leaves and crowns symptoms were captured by at 7 or 45 days post inoculation, respectively, and used for disease severity analysis as described in prerious studies (Wang et al., 2023; Wei et al., 2024). The population of Xaf YL19-GFP on strawberry leaves were collected at 7 dpi and were measured using RT-qPCR as described previously (Wang et al., 2023) .

### 3.4 Immunoprecipitation and protein analysis

Strawberry or *N.benthamiana* plant leaves were frozen and crushed in liquid nitrogen to determine the protein content. The fine sample powder was boiled in water for five minutes in SDS loading buffer, and the supernatant was utilized for immunoblot analysis after the sample was centrifuged for ten minutes at 12,000 g. Before collecting samples and freezing them in liquid nitrogen, *N. benthamiana* leaves were co-infiltrated with agrobacterium carrying XopL-GFP and FveACO9-Myc or free GFP and FveACO9-Myc for expression. This allowed for the verification of effector-target interaction. The Co-immunoprecipitation (Co-IP) experiment was performed in accordance with Xian et al. (2020) guidelines. To immune-precipitate FveACO9-GFP or free GFP, the leaf lysate from the co-infiltrated leaves of *N. benthamiana* with 20 µl of GFP-Trap beads (ChromoTek, Germany) was incubated at 4 °C for 2 hours. The beads were washed five times with PBS solution before being loaded with 50 µl SDS sample loading buffer and 20 µl boiling protein samples for immunoblotting and SDS-PAGE gels. Each immunoblot was assessed using the appropriate antibodies anti-GFP, anti-Myc, or anti-luciferase (Biodragon, China). Blots were stained with Ponceau S to verify equal loading.

### 3.5 RNA isolation and quantitative RT-PCR

Total RNA was isolated from strawberry leaves (50 mg) using the E.Z.N.A.^®^ Total RNA extraction kit II according to the manufacturer’s protocol (Omega, BIO-TEK, USA). Complementary DNA (cDNA) was synthesized with TransScript^®^ One-Step gDNA Removal and cDNA Synthesis SuperMix (TransGen, Beijing, China) using total RNA as a template. Quantitative RT-PCR (qRT-PCR) was performed using an Applied Biosystems^TM^ StepOnePlus^TM^ Real-Time PCR System (ThermoFisher, USA). The gene expression levels were calculated based on three repeats and were normalized against the expression of *FveGAPDH* (NCBI GenBank ID: XM_004287072.2). The primers used for qRT-PCR were listed in Supplementary Table 3.

### 3.6 Plant ethylene inhibitor or promoter pretreatments

Before inoculation, plants were subjected to different pretreatments with 30 μM ACC, 50 μM AIB (α-aminoisobutyric acid, the ACC oxidases inhibitor), and the control plants were applied with H_2_O. The ethylene level of each treatment was measured to confirm the effect of ethylene inhibition or promotion 12 hours after treatment, then plants were inoculated with *Xaf* YL19 and measured for bacteria population as described above.

### 3.7 Yeast two-hybrid (Y2H) and Y2H-sequence assays

For the Y2H assay, the coding region of effectors was inserted into the pGBKT7 vector (BD) or pBT3-N to make the construct pGBKT7-T3E or pBT3-N-T3E, which was subsequently transformed into the yeast Y2H-Gold or NMY51 with the lithium acetate transformation method, respectively. After confirming that effectors or their derivates do not have autoactivation activity in Y2H-Gold or NMY51 (Fig S1), T3Es were used as the bait to screen target proteins from the mixed cDNA library of *F.vesca* ‘Hawaii 4’ leaves challenged with *X.fragariae* YL19. The coding region of each candidate target was cloned from *F.vesca* ‘Hawaii 4’ cDNA and introduced into pGADT7 vector (AD) or pPR3-N for interaction verification by co-transformation of the prey and the bait to Y2H-Gold or NMY51, respectively. Transformants were placed on SD-Leu-Trp (double dropout, DDO) plates and incubated for 2 d at 30 ℃. The interactions were tested on SD-Trp-Leu-His-Ade (quadruple dropout, QDO) plates and incubated for 5-7 d at 30℃. All freshly grown yeast colonies grown on QDO plates were scratched off, and plasmid DNA was prepared from the yeast pellet using a standard alkaline lysis protocol and ethanol precipitation. The samples were sequenced by Illumina Hi-Seq 2000 125-bp paired-end reads as described by Chen et al. (2021)

### 3.8 Luciferase-based protein stabilization assay

The firefly luciferase (LUC) coding region was tagged to the C-terminal of FveACO9 and co-expressed with XopL-GFP in *N.benthamiana*, GFP/FveACO9-LUC, XopL-GFP/LUC, and GFP/LUC co-expressed in *N.benthamiana* as controls. The LUC images were captured and luminescence intensity was quantified on the infiltrated leaves 72 hours post agroinfiltration immediately after spraying luciferin using Plant view 100 cooled CCD imaging apparatus (MTL, China). The protein levels of XopL-GFP, GFP, and FveACO9-LUC were detected by immunoblotting with corresponding anti-GFP or anti-luciferase (Biodragon, China, 1:5, 000) monoclonal antibodies.

### 3.9 ACO enzyme activity assay

The total ACO enzyme activity of *XopL*-OX strawberry lines was measured based on the method of Bulens et al. (2011). 0.5 g strawberry leaves were ground in 1 ml of MOPS extraction buffer (400 mM MOPS, 30 mM ascorbic acid sodium salt, and 10% (v/v) glycerol) at pH 7.2, and gentle shaking incubation at 4℃ for 20min. The supernatant was separated for analysis by centrifuging it at 14000 g for 20 minutes at 4℃. 400µl of strawberry extract, 50 mM MOPS, 150 mM NaHCO_3_, 30 mM ascorbic acid, 1.0 mM ACC, 0.1 mM FeSO_4_, 10% (v/v) glycerol, 0.1 mM DTT, and a 10 mL headspace container held a 3.6 mL mixture. One hour of shaking incubation at 30℃ resulted in the extraction of 1 mL of gas for ethylene measurement using gas chromatography (Agilent 7820A, USA).

### 3.10 Statistical analysis

The data in each figure are from representative experiments that were independently repeated at least three times. Statistical analyses were performed with a two-sided *t*-test, or one-way ANOVA, following multiple comparisons of means with Duncan’s multiple range test. Figures were drawn using the GraphPad Prism 9 software (San Diego, USA).

## Accession numbers

The accession numbers for the genes in this study are as follows: *FvePR1* (pathogenesis-related protein-1-like, FvH4_2g02920.1), *FvPR5.3* (Thaumatin-like protein, FvH4_6g16950.1).

## Credit authorship contribution statement

Xiao-Lin Cai: Investigation, Data curation, Methodology, Formal analysis, Writing – original draft, review & editing. Wenyao Zhang: Investigation, Methodology, Formal analysis–review & editing. Jinnan Luo: Formal analysis–review & editing. Wei Li: Investigation. Ruihong Chen: Investigation. Ying-Qiang Wen: Conceptualization, Project administration, Funding acquisition. Jia-Yue Feng: Conceptualization, Project administration, Funding acquisition.

## Acknowledgments

The authors would like to thank the anonymous reviewers for their comments on the manuscript. This work was sponsored by the Key Technology Tackling Project of Agricultural Key Industry Chain in Xi’an (No. 22NYGG0006), major project of the Agricultural Cooperative Innovation and Extension Alliance in Shaanxi Province (No. LMZD202102), the Key Research and Development Program of Shaanxi province (No. 2023-YBNY-083), and the Scientific and technological innovation and achievement transformation project of experimental demonstration station (No. TGZX2021-20).

